# CYTOTOXIC ACTIVITY OF A NEW ISOFORM L-AMINO ACID OXIDASE (BALT-LAAO-II) FROM *Bothrops alternatus* (URUTU) SNAKE VENOM IN HUMAN LEUKEMIC HL60 CELLS

**DOI:** 10.1101/2020.01.15.907758

**Authors:** Maurício Aurelio Gomes Heleno, Alexandre Nowill, João Ernesto de Carvalho, Diego L. Suni-Curasi, Julissa Vilca-Quispe, Emilio Alberto Ponce-Fuentes, Gustavo Alberto Obando-Pereda, Luis Alberto Ponce-Soto

**Author notes:** corresponding author: Prof. Luis Alberto Ponce-Soto Ph.D., Laboratorio de Quimica de Proteinas, Universidad Catolica de Santa Maria, Arequipa - PERU, anexo 1601.

## Abstract

In this work we describe the isolation of a new isoform L-amino acid oxidase (LAAO) referred to as Balt-LAAO-II from *Bothrops alternatus* snake venom, which was highly purified using a combination of molecular exclusion (Sephadex G-75) and RP-HPLC chromatographics steps. When analyzed by SDS-PAGE, the purified Balt-LAAO-II presented a molecular weight of ∼66 kDa. The N-terminal amino acid sequence and internal peptide sequences showed close structural homology to other snake venom L-amino acid oxidases.

This enzyme induces *in vitro* cytotoxicity on cultured human leukemic HL60 cells. Cells were grown in RPMI medium and were incubated with isoform Balt-LAAO-II (1, 10 and 100 μg/mL) for up to 72 h. All three concentrations of venom markedly decreased the cell viability from 6 h onwards based on the staining with propidium iodide, the reduction of 3-(4,5-dimethylthazol-2-yl)-2,5-diphenyl tetrazolium bromide (MTT) and the uptake of neutral red.

Flow cytometry showed that all isoform Balt-LAAO-II and whole venom concentrations induced apoptosis after 2-6 h of incubation. Morphological analysis of cells incubated with isoform Balt-LAAO-II and whole venom showed cell rounding and lysis that increased with the venom concentration and duration of incubation. These results show that isoform Balt-LAAO-II from venom *Bothrops alternatus* is cytotoxic to cultured HL60 cells and suggest that this damage may involve apoptotic and oxidative stress pathways.

## 1. Introduction

The snake *Bothrops alternatus* occurs in southeastern and southern Brazil, Paraguay, Uruguay and northern Argentina (Campbell and Lamar, 1989). The venom of this species contains enzymes common to other *Bothrops* spp., including a thrombin-like enzyme (Smolka *et* al., 1998), a non-enzymatic thrombin inhibitor (Castro *et al*., 1998), metalloproteinases/ disintegrins (Souza *et al*., 2000; Cominetti *et* al., 2003), phosphodiesterase (Valério *et al*., 2002) and phospholipase A_2_ (Nisenbom *et al*., 1986a,b, 1988). One isoform of L-amino acid oxidase, descried as Balt-LAAO-I is an acidic glycoprotein, pI 5.37, homodimeric, Mr 123, 000, whose N-terminal sequence is ADVRNPLE EFRETDYEVL … and induces platelet aggregation and shows bactericidal activity against *Escherichia coli* and *Staphylococcus aureus*. In addition, this enzyme is slightly hemorrhagic and induces edema in the mouse paw (Rodrigo G. Stábeli *et* al., 2004) and PLA_2_ homólogous K49 (Ponce-Soto *et* al., 2007, 2009). Although *Bothrops alternatus* venom can produce local cellular damage such as necrosis, little is known about its cytotoxic action on malignant cells.

L-Amino acid oxidases (LAAOs, EC 1.4.3.2) are flavoenzymes catalyzing the stereospecific oxidative deamination of a wide range of L-amino acids to form corresponding α-keto acids, H_2_O_2_ and ammonia via an imino acid intermediate. LAAOs are widely distributed in the venomous snake families Viperidae, Crotalidae and Elapidae (Du and Clemetson, 2002; Tan and Fung, 2009). The enzyme exhibits a wide range of biological activities including apoptosis-inducing, edema-inducing, inhibition or induction of platelet aggregation, antibacterial effect and antiviral activity (Tan and Fung, 2009). They can also have antimicrobial, antiprotozoal or anticoagulant activity, be cytotoxic and antiproliferative on tumour cells (Santos *et* al., 2004, Izidoro, et al., 2006). The role of LAAO in the pharmacological action of snake venom, however, is still not fully understood. Generally, the enzyme has a low lethal toxicity in mice.

Apoptotic processes and cellular damage are mechanisms of action of some toxins, and several studies show potential applications of such substances as models for the development of chemotherapy and anticancer agents. In view of the lack of information, we have purified an isoform of L-amino acid oxidase (LAAO) from *Bothrops alternatus* venom named Balt-LAAO-II and have studied some of its biochemical and cytotoxicity activity on cancer cells of the human leukemic cell line HL60. Our results have important implications for understanding the envenomation by *Bothrops alternatus* as well as the mechanisms of cell death triggered by this group of enzymes and provide a foundation for further investigations.

## 2. Material and Methods

### 2.1 Venom and reagents

*Bothrops alternatus* venom was a gift from the Batatais Serpentarium, Batatais, SP, Brazil. All chemicals and reagents used in this work were of analytical or sequencing grade. Fetal calf serum (FCS), 3-(4,5-dimethylthazol-2-yl)-2,5-diphenyl tetrazolium bromide (MTT), glutamine, sodium pyruvate, penicillin and streptomycin were obtained from Sigma Chemical Co. (St. Louis, MO, USA), aracityn was from Pfizer, and RPMI medium was from Gibco Life Technologies (Paisley, Scotland). Annexin V/propidium iodide apoptosis detection kits were obtained from R&D Systems Inc. (Minneapolis, MN, USA). Sterile multiwell culture plates were from Corning Inc. (Corning, NY, USA). The other reagents of analytical grade were from Baker, Mallinkrodt or Merck.

### 2.2 Isolation of of L-amino acid oxidase from Bothrops alternatus venom

Molecular exclusion chromatography: Approximately 35 mg of the whole venom of *Bothrops alternatus* was dissolved in 400μl ammonium bicarbonate buffer (0,2M; pH 8.0) and homogenized up to complete dissolution, followed by clarification with high speed centrifugation (4500x *g* for 2min). The supernatant was fractionated on a Sephadex G75 (Pharmacia) column (1.6 × 100cm) pre-equilibrated with ammonium bicarbonate buffer. The flow rate was 0.2 ml/min and the elution profile was monitored at 280 nm. The fractions collected were immediately lyophilized and stored at -40°C.

The samples were applied at a flow rate of 1 mL/min onto reversed-phase liquid chromatography (RP-HPLC) C18 column (20 × 250 mm). Fractions containing LAAO were pooled and stored at -20°C.

The molecular homogeneity of this protein was evaluated by reverse-phase HPLC (0.1 × 30 cm column of µ-Bondapack C-4, Waters) using a linear and discontinuous gradient 0– 100% of acetonitrile in 0.1% trifluoroacetic acid (TFA) (v/v). Afterwards, 3 mg of the LAO protein from the RP-HPLC was dissolved in 250 mL of buffer A (0.1% TFA) and centrifuged at 4500g for 2 min and the supernatant was then applied on the analytical reverse-phase HPLC, previously equilibrated with buffer A for 15 min. The elution of the protein was then conducted using a linear gradient of buffer B (66.6% acetonitrile in buffer A) monitored at 280nm. After elution, the fraction was lyophilized and stored at -40°C. The purity of the LAO was also assayed using PAGE-SDS electrophoresis.

### 2.3 Protein determination and L-amino acid oxidase assay

Total protein was evaluated by the microbiuret assay (Johnson, 1978). L-amino acid oxidase (15 μg) activity was determined by a spectrophotometric assay using L-leucine as substrate (Pessatti et al., 1995). In this assay, the oxidative deamination of L-leucine produced hydrogen peroxide, which was reduced in the presence of horseradish peroxidase (Biolabs-INC, New England) by *o*-phenylenediamine to produce a colored oxidized product, which was spectrophotometrically monitored at 490 nm. The assay mixture with 10 mL of L-leucine 1% in Tris–HCl 0.1 M (pH 7.2) contained 16 μL horseradish peroxidase (1 mg/mL) and 100 μL *o*-phenylenediamine (10 mg/mL in methanol). After 30 min, the reaction was stopped by addition of 10% (w/v) citric acid. For enzymatic specificity assays, L-leucine was replaced by other L-amino acids (2 mmol), namely: Leu, Ile, Met, Cys, Val, Tyr, Trp, Gln, Thr, Ser, Lys, Arg, Phe, Asn, Glu, Gly, Pro, Asp and His, at the same concentration and assayed for activity under identical conditions.

### 2.4 Cell culture

HL60 cells were grown in RPMI medium supplemented with 10% FCS and 2 mM L-glutamine, 1 mM sodium pyruvate, 100 μg of penicillin/mL and 100 μg of streptomycin/mL in a humidified atmosphere with 5% CO_2_ at 37^°^C. The cytotoxicity of the venom was tested by incubating the cells with medium containing *Bothrops alternatus* venom (1, 10 and 100 μg/mL) or fractions (10µg/mL) for 2, 6, 24, 48 and 72 h.

### 2.5. Cytotoxicity assays

#### 2.5.1 Neutral red uptake

Cells grown in 96-well plates were treated with LAAO venom as described above. After incubation with venom the cells were centrifuged and the was medium removed from the plates prior to the addition of 200 μl of neutral red solution (Seromed; final concentration of 50 μg/mL) were added to each well. The plates were then incubated for 3 h at 37^°^C, after which the cells were centrifuged and the medium was removed. The cells were then rapidly washed with 200 μL of fixative (1% v/v formaldehyde, 1% w/v CaCl_2_), centrifuged and the fixative removed. One hundred microliters of a solution of 1% acetic acid and 50% ethanol was then added to the wells to extract the dye. After incubation for 1 h on a microtiter plate shaker at room temperature, the plates were read at 540 nm on a microplate reader (SpectraMax 340, Molecular Devices, Sunnyvale, CA, USA).

#### 2.5.2 MTT reduction

Cells grown in 96-well plates were treated with LAAO as described above. At the end of the incubations, the cells were centrifuged, the supernatant was removed and 10 μL of a solution of MTT (3-(4,5-dimethylthazol-2-yl)-2,5-diphenyl tetrazolium bromide) (5 mg/mL) and 100 μl of PBS were added to each well. After incubation for 3 h at 37^°^ C, 100 μL of isopropanol were added to the wells and the plates then incubated with shaking for 10 min prior to reading the absorbance at 570 nm in a SpectraMax microplate reader

#### 2.5.3 Cell viability and flow cytometry

Cell viability was assessed by staining with propidium iodide, essentially as described below for flow cytometry. HL60 cells control, annexin, whole venom from *Bothrops alternatus* and Balt-LAAO-II (10 μg/mL). Cells grown in six-well plates were treated annexin, whole venom from *Bothrops alternatus* and Balt-LAAO-II as described above and processed for the detection of apoptosis by flow cytometry using a commercial kit. At the end of each incubation with venom, the cells were pelleted by centrifugation (200 x *g*, 10 min, ∼25^°^C), the supernatant was removed and 2 ml of PBS was added. After a further centrifugation, the supernatant was discarded and 400 µl of cold binding buffer (0.1 M Hepes/NaOH, pH 7.4, 1.4 M NaCl, 25 mM CaCl_2_) were added to each tube followed by vortex mixing and incubation for 15 min on ice. After this period, 10 μl of concentrated calcium solution was added, and the tubes vortexed prior to the addition of 1 μL of annexin V to each tube. The tubes were vortexed again and 10 μL of propidium iodide were added to the tubes. The cells were analyzed by flow cytometry (BD Technologies) immediately or stored on ice in the dark until analysis (within 30 min).

### 2.6 Morphological observations

The changes in cell appearance following incubation with whole venom from *Bothrops alternatus* and Balt-LAAO-II, were monitored with a Zeiss inverted light microscope fitted with a digital camera. The venom-induced alterations were compared with corresponding control cells.

### 2.7 Statistical analysis

The results were expressed as the mean±S.E.M., as appropriate. Statistical comparisons between groups were done using analysis of variance (ANOVA) followed by the Bonferroni test. Values of *p*<0.05 were considered significant.

## 3. Results

### 3.1 Biochemical and structural analysis of L-amino acid oxidase (Balt-LAAO-II) from Bothrops alternatus (urutu) snake venom

The fractionation of *Bothrops alternatus* snake venom on Sephadex G-75 column allowed the purification of three major fractions, named fractions I, II, III, IV and V (Fig. 1). Enzymatic assay of fraction I confirmed the characteristic action described for L-amino acid oxidase (LAAO).

**Figure 1.**
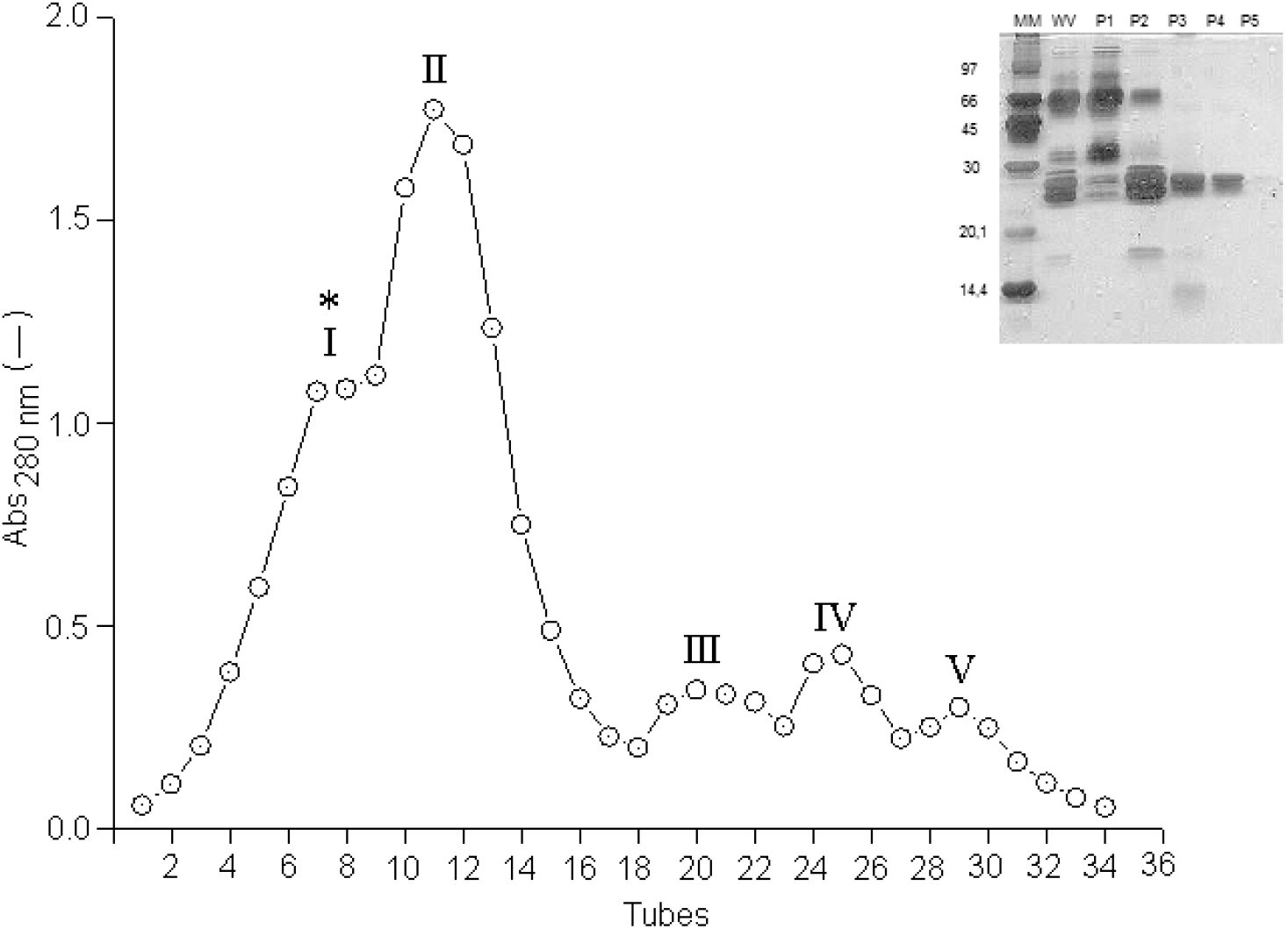
Molecular exclusion chromatography of *Bothrops alternatus* snake venom on a Sephadex G-75 column (1.6 cm x 100 cm) at 14^°^C. 35 mg of crude venom were applied to the column, equilibrated with ammonium bicarbonate buffer 0,2M; pH 8.0 at a flow rate of 0.3 ml/min. The elution profile was monitored at 280 nm. The main fractions obtained identified as I-V (L-amino acid oxidase peak I.) were pooled, lyophilized, and stored at -20°C. Insert: SDS-PAGE main fractions obtained by molecular exclusion chromatography.

Re-chromatography of L-amino acid oxidase (fraction I) by ion exchange HPLC Protein Pack DEAE 5PW column, produced nine peaks, the main ones being I-1 – I-9 (Fig. 2). All fractions obtained by this procedure were eluted with less than 0.5 M buffer and were easily lyophilized with no need for dialysis. The peaks I-3 and I-4 showed isoforms of L-amino acid oxidase.

**Figure 2.**
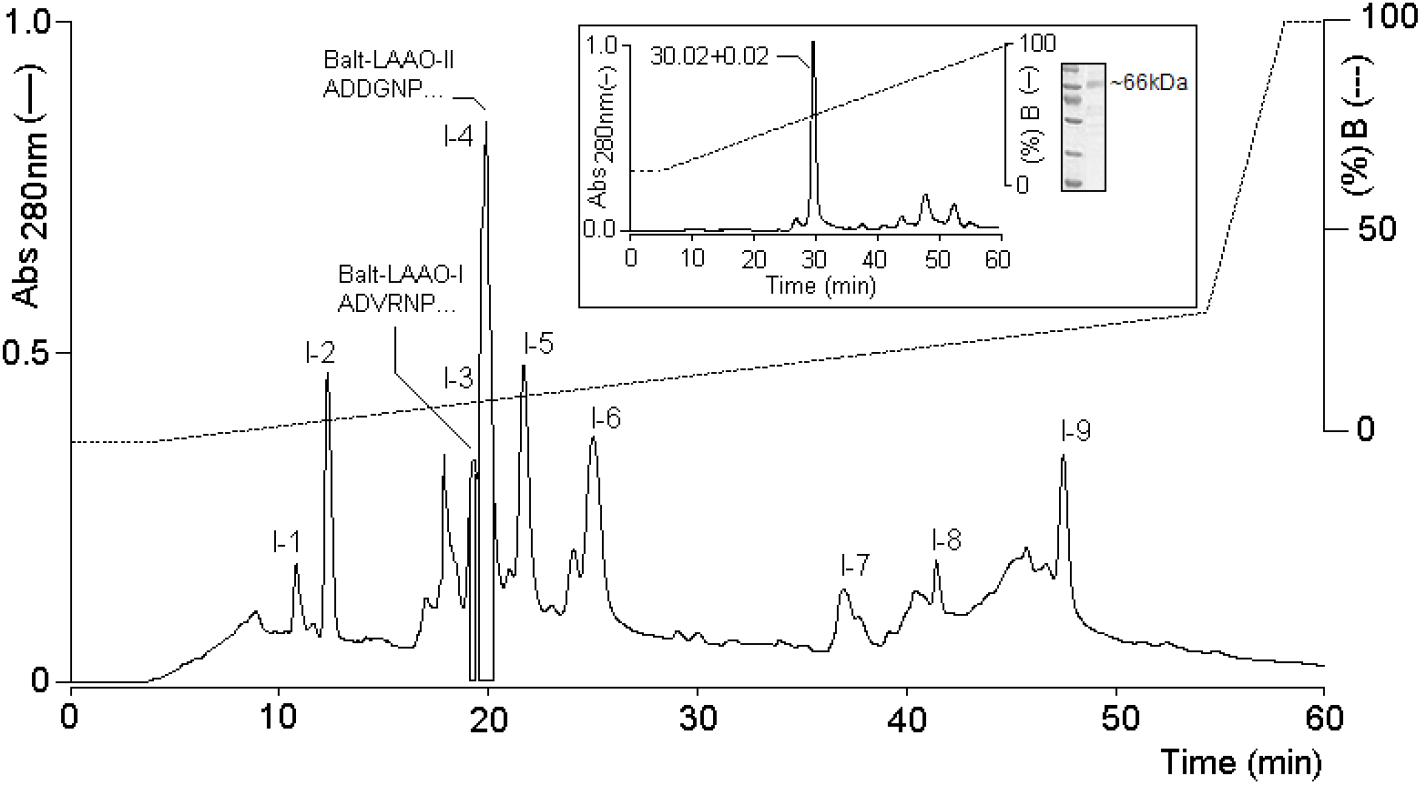
Chromatography on RP-HPLC of the peak I from molecular exclusion chromatography of lyophilized from *Bothrops alternatus* snake venom. The main fractions obtained identified as I-1 – I-9 (L-amino acid oxidase peak I-4, Balt-LAAO-II.) was pooled, lyophilized, and stored at -20°C. Insert: Re-chromatography and SDS-PAGE of main fraction (Balt-LAAO-II) obtained by RP-HPLC.

The re-chromatography of peak I-4 by RP-HPLC resulted in one peaks, with retention times corresponding to 30.2 ± 0.02 min (I-4) of time retention. (Fig. 2 insert). The N-terminal sequences of isoform I-4 (Balt-LAAO-II) showed high amino acid sequence identity with other LAOs (Fig. 3).

**Figure 3.**
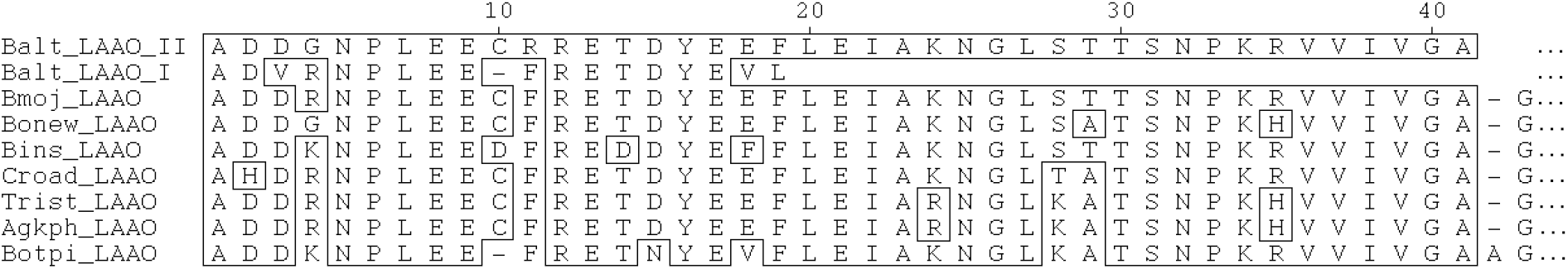
N-terminal amino acid sequence of L-amino acid oxidase from the *Bothrops alternatus* venom (Balt-LAAO-II) compared with others LAAO, obtained from the BLAST similarity search (PubMed, MEDLINE): Bmoj_LAAO (*Bothrops moojeni*, Q6TGQ8), Bonew_LAAO (*Bothrops neuwiedi*), Croad_LAAO (*Crotalus adamanteus*, 093364), Bins_LAAO (*Bothrops insularis*, Braga et al., 2008), Trist_LAAO (*Trimeresurus stejnegeri*, Q6WP39), Agkph_LAAO (*Agkistrodon piscivorus*; Q6STF1) and Botpi_LAAO (*Bothrops pirajai*, Izidoro et al., 2006)

The first 40 amino acid residues of Balt-LAAO-II were identified by automated Edman degradation and provide additional evidence for the purity of the enzyme and homology between different venom L-amino acid oxidases (Fig. 3). Comparison of the N-terminal sequence of Balt-LAAO-II with that of enzymes isolated from other snake venoms revealed high homology with LAAOs from viperid venoms. In the N-terminal sequence, at least 20 of the 40 amino acid residues were found to be fully conserved in all sequences, thus suggesting the presence of a highly conserved Glu-rich motif.

The affinity of Balt-LAAO-II to different substrates was accessed for further biochemical characterization. The enzyme showed higher affinity to the hydrophobic amino acids L-Met, L-Leu, L-Phe and L-Ile. For other amino acids, the catalytic affinity was very low (Ala, Val, Glu and Tyr) or inexistent (L-Pro, L-Thr, L-Ser and L-Cys) (Data not show).

### 3.2 Cytotoxicity assays

Figure 4, shows the cell viability assessed by staining with MMT reduction (A) after incubation with isoform L-amino acid oxidase Balt-LAAO-II for various periods of time. All L-amino acid oxidase Balt-LAAO-II concentrations caused a decrease in cell viability after 6 h that was very marked after 24 h. The decrease in Neutral red (B) and Blue trypan uptake (C) caused by L-amino acid oxidase Balt-LAAO-II paralleled the changes in viability.

**Figure 4.**
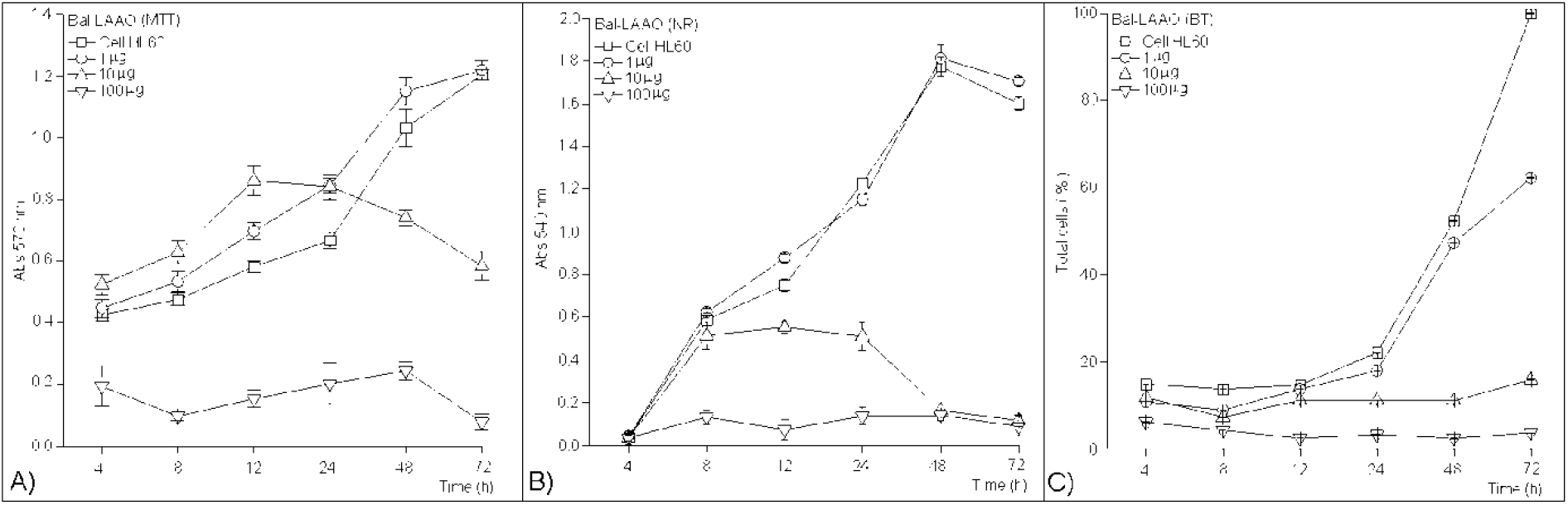
(A) Cell viability MTT reduction, (B) Neutral red and (C) Blue trypan uptake by HL60 cells incubated with different concentrations of Balt-LAAO-II isolated from *Bothrops alternatus* snake venom for the indicated times. The columns are the mean + 1 S.E.M. of 5 experiments. \**p*<0.05 compared to the corresponding control.

### 3.3 Venom-induced apoptosis

Flow cytometry of HL60 showed that whole venom and L-amino acid oxidase Balt-LAAO-II (10 μg/mL) from *Bothrops alternatus*, caused apoptosis from 6h onwards that was maximal after 24 h (Figure 5C and D. Figure 5A and B control negative and positive). Number of cells showing apoptosis after incubation with different concentrations of fractions obtained by gel filtration Sephadex G-75 from *Bothrops alternatus* snake venom (10 µg/mL) and (B) differents concentrations of Balt-LAAO-II. Aracityn (20 nM) was used as a positive control (Figure 6).

**Figure 5.**
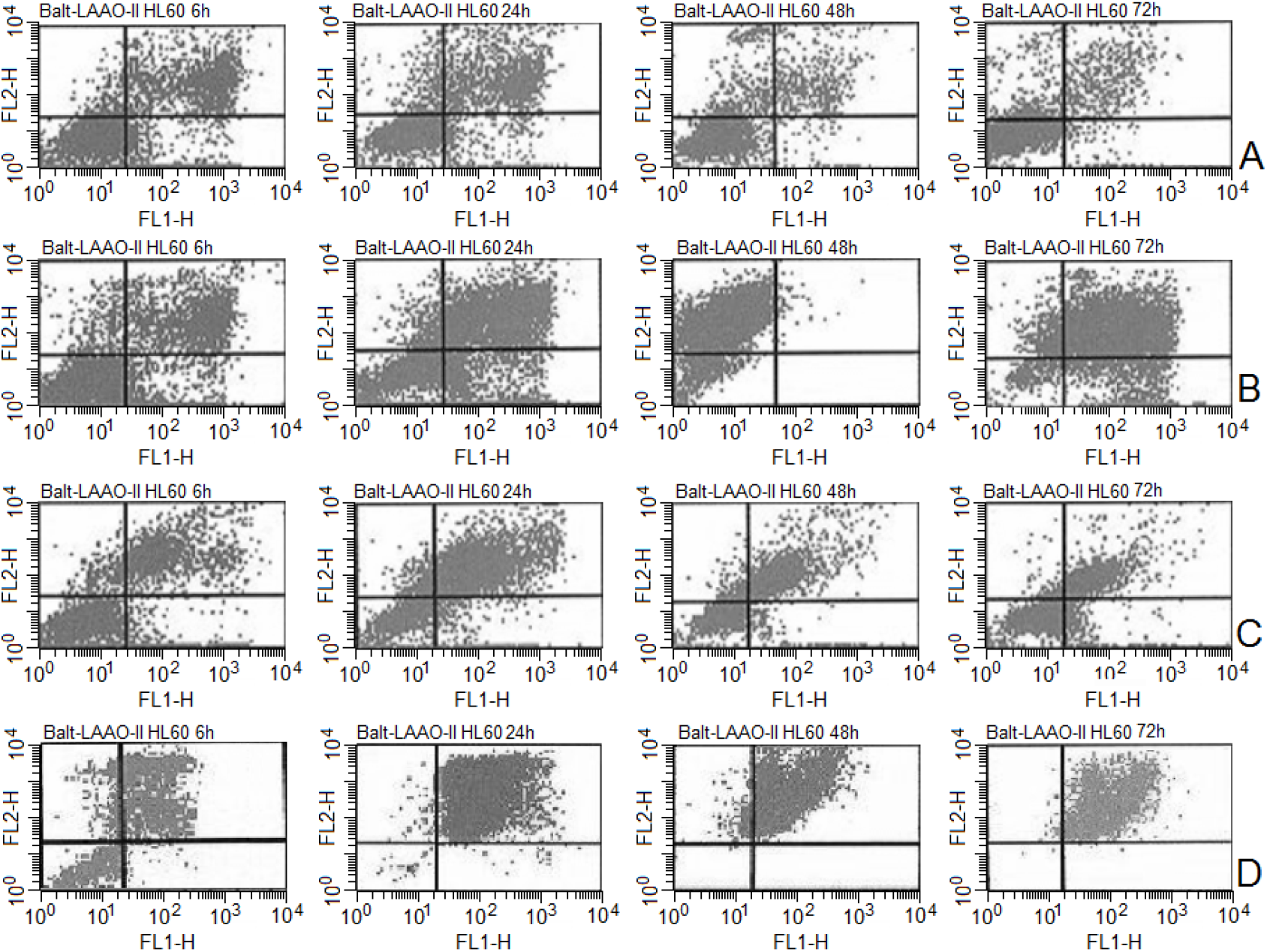
Flow cytometer scattergrams showing apoptosis in HL60 cells control (A), treated with aracityn (B) (10 μg/mL), wolhe venom (10 μg/mL) (C) and Balt-LAAO-II (10 μg/mL) (D) for 6, 24, 48 and 72 h, respectively.

**Figure 6.**
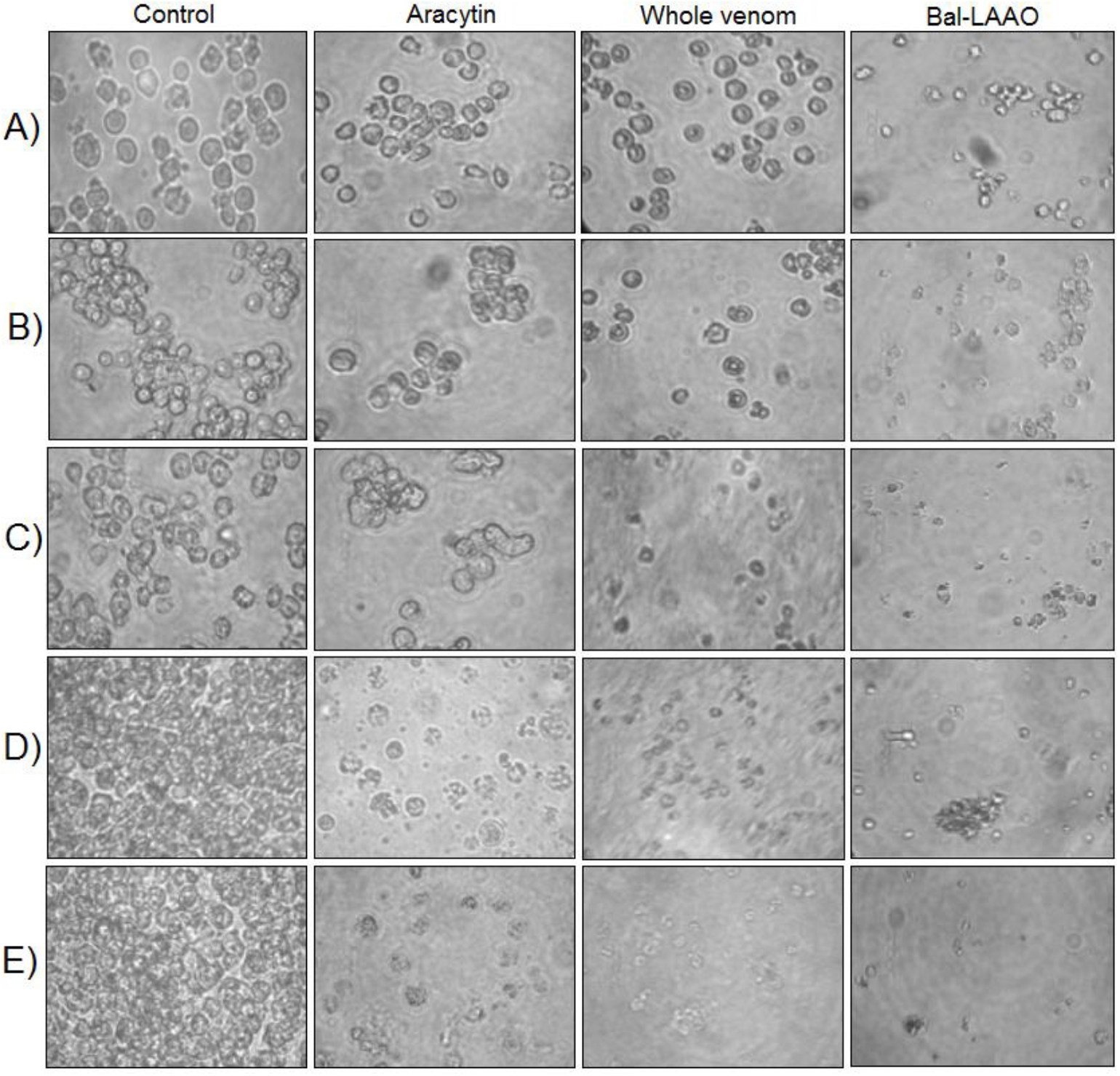
(A) Number of cells showing apoptosis after incubation with different concentrations of fractions obtained by gel filtration Sephadex G-75 from *Bothrops alternatus* snake venom (10 μg/mL) and (B) differents concentrations of Balt-LAAO-II. Aracityn (20 nM) was used as a positive control.

### 3.4 Cell morphology

Figure 7 shows that the incubation of HL60 cells with venom *Bothrops alternatus* and L-amino acid oxidase Balt-LAAO-II, produced marked time-dependent changes in cell morphology, with the main alterations being rounding of the cells and subsequent lysis.

**Figure 7.**
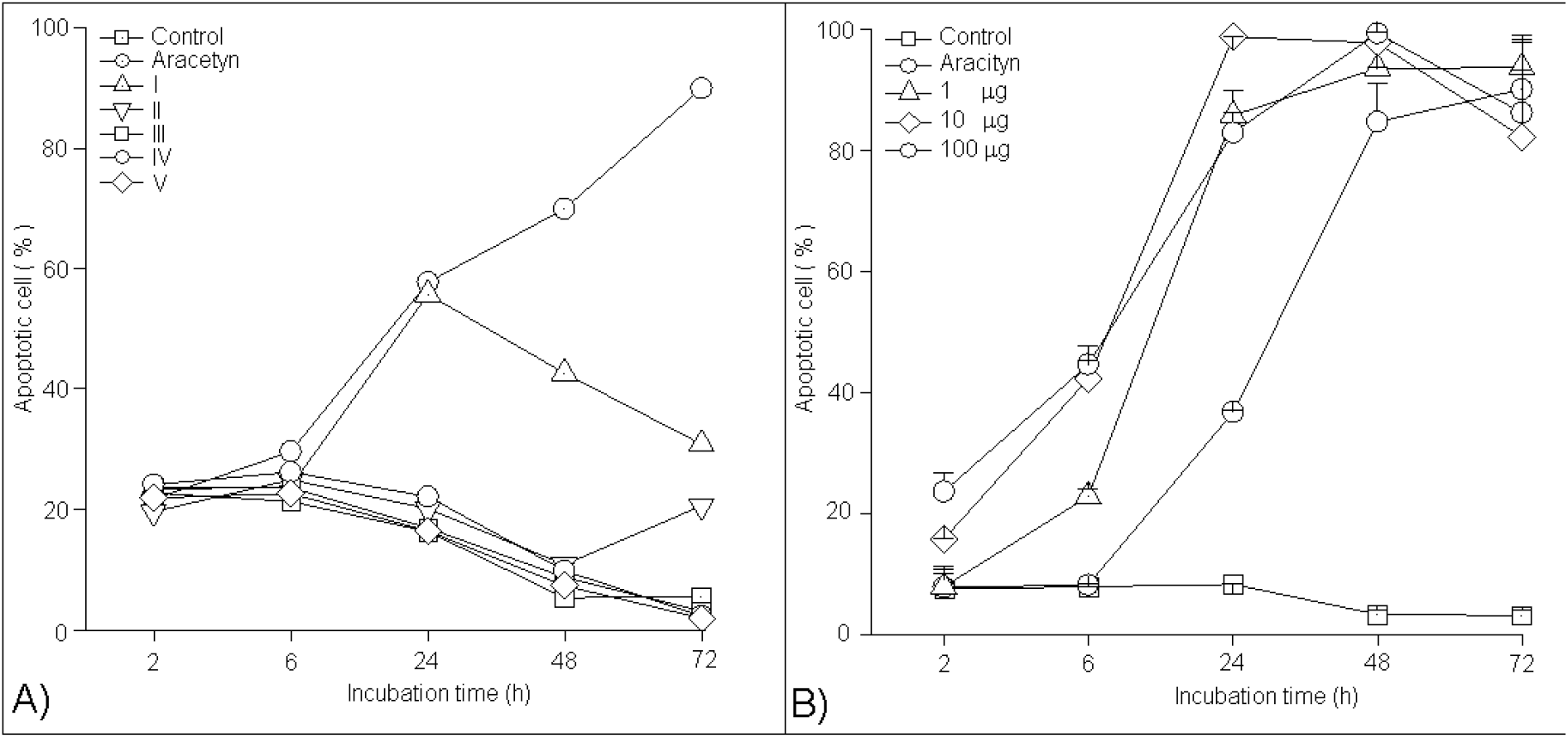
Morphological alterations in HL60 cells treated with *Bothrops alternatus* venom (100 μg/ml) and L-amino acid oxidase Balt-LAAO-II (10 μg/mL) for 2, 6, 24, 48 and 72 h (A, B, C, D and E, respectively. Aracityn (20 nM) was used as a positive control.

## 4. Discussion

L-Amino acid oxidases are widely expressed in snake venoms and catalyze the oxidative deamination of L-amino acids, producing the corresponding α-ketoacids, hydrogen peroxide and ammonia. They are usually homodimeric, FAD-binding glycoproteins, with molecular mass around 110–150 kDa when measured by gel filtration under non-denaturing conditions. However, the molecular mass of L-Amino acid oxidases from snake is around 50–70 kDa when assayed by SDSPAGE, both under reducing and non-reducing conditions. Thus, most of the L-Amino acid oxidases from snake are homodimeric glycoproteins associated by non-covalent bonds with pI of approximately 4.4–8.2 (D. Butzke et al., 2005, G. Ponnudurai et al., 1994, Y. Jin et al., 2007, Raquel de Melo Alves Paiva et al., 2011. Gustavo B. Naumann et al., 2011).

In the present work, we isolated a new isoform of L-Amino acid oxidases from *Bothrops alternatus* snake venom (Balt-LAAO-II) by using two chromatographic steps (molecular exclusion in Sephadex G75 and ion exchange Protein Pack DEAE 5PW column HPLC) and finally the active fraction was still analyzed for purity by reverse phase HPLC chromatography. Balt-LAAO-II presented molecular weight of approximately ∼66 kDa in SDS-PAGE.

The enzyme showed high affinity for L-Leu is the most favorable substrate for evaluating the enzymatic activity of purified snake venom L-Amino acid oxidases, but high catalytic activity was detected against L-Met for Balt-LAAO-II. At the same way, L-Amino acid oxidases from *Bothrops alternatus, Bothrops pirajai* and *Bothrops moojeni* favor the specific substrates of hydrophobic amino acids (L-PheNL-MetNL-LeuNL-Ile) (França et al., 2007; Stábeli et al., 2004). The catalytic differences may be explained by differences in side chain binding sites responsible for the substrate specificity of the enzyme (Samel et al., 2006).

The N-terminal sequence of amino acids shows a high homology to other members of this family of toxins, including the presence of a highly conserved Glu rich motif, which may play a role in substrate binding. In agreement with the structure of L-Amino acid oxidases from *Calloselasma rhodostoma* (P. Macheroux et al., 2001), this N-terminal region (5–25 residues) constitutes one part of the substrate-binding domain. Our finding of 2-DE that the enzyme has a range of isoelectric points (pI) from about 5.8 to 6.1, reflects the presence of isoforms with distinct pIs similarly to other snake venom L-Amino acid oxidases (Y.H. Sakurai et al., 2001, S.A. Ali et al., 2000, P.D. Pawelek et al., 2000). However, the existence and the nature of the possible isoforms have to be further investigated.

The presence of isoforms may be due to different composition or different glycosylation as revealed for other homologous flavoenzymes (Sakurai et al., 2001, R.G. Stabeli et al., 2004), or they may have been synthesized from different genes as was suggested for two distinct L-Amino acid oxidases from *Pseudechis australis* (B.G. Stiles et al., 1991). Moreover, heterogeneous glycosylation of isoform Balt-LAAO-II may also cause this divergence and would not be surprising as the venom used for purification of Balt-LAAO-II was collected from only two snake individuals.

Venomous animals have evolved a vast array of toxins for prey capture and defense. Toxins as L-amino acid oxidases have been reported to show potent applications in pharmacology and cancer therapy (X.Y. Du, K.J. Clemetson 2002). As the balance between therapeutic potential and toxic side effects of a toxin is very important when evaluating its usefulness as a pharmacological drug, experiments were designed to investigate the in vitro cytotoxicity of Balt-LAAO-II against human cancer cells (HL-60).

Apoxin-I, a *Crotalus atrox* L-amino acid oxidases, induced apoptosis mediated by H_2_O_2_ S. (Torii et al., 2000). However, Suhr et al., (1996) demonstrated that the L-amino acid oxidases induced apoptotic mechanism is clearly distinguishable from the one stimulated directly by exogenous H_2_O_2_, suggesting that the L-amino acid oxidases induced apoptosis was not solely due to the H_2_O_2_ produced by the enzymatic reaction. However, the molecular details as to which intracellular components are specifically associated with L-amino acid oxidases induced apoptosis are still unknown. In addition, Balt-LAAO-II induced mouse paw edema (data not shown). However, the enzyme is neither hemorrhagic nor myotoxic at doses of up to 100 µg/animal (data not shown). The physiological role of L-amino acid oxidases in snake envenomation is not well understood. Venom L-amino acid oxidases share many biological effects, and this study and others demonstrated that these effects were mainly due to the production of H_2_O_2_ by the enzymatic reaction. The exact molecular mechanisms of these effects are still being investigated. Balt-LAAO-II induced dose-dependent cytotoxicity in human leukemic HL60 cells.

The close correlation between cell viability and the decrease in the uptake of neutral red, blue trypan and in MTT reduction, which assess lysosomal and mitochondrial function, respectively, suggested that the cytotoxic action of the venom involved primarily cell lysis rather than the disturbance of energy production and metabolism (Figure 4A, B and C). Other studies have also reported an antiproliferative action for *Bothrops* venoms and their components both *in vitro* and *in vivo* (Pereira-Bittencourt *et al*., 1999; de Carvalho *et al*., 2001; Corrêa *et al*., 2002; da Silva *et al*., 2002a, b).

Balt-LAAO-II and whole venom from *Bothrops alternatus* snake venom, induced cell death in HL-60 cell line. In this study, flow cytometry (Figure 5) was utilized to confirm the results obtained from morphological assessment of cell death by microscopy (Figure 6) The morphological alterations seen in HL60 cells following incubation with venom (rounding and lysis) were similar to those reported for other *Bothrops* venoms (Collares-Buzato *et al*., 2002) and their isolated components (Borkow *et al*., 1995; Corrêa *et al*., 2002). In adherent cells, loss of contact with the substrate partly reflects the action of venom disintegrins which bind to cell adhesion molecules (integrins) present on the cell surface (Kamiguti *et al*., 1998). The integrins known to be affected by *Bothrops* venom disintegrins include α1β3 (Rucinski *et al*., 1990), α2β1 (Souza *et al*., 2000; Moura-da-Silva *et al*., 2001) and α5β1 (Cominetti *et al*., 2003). In contrast, αIIβ3, α1β1, α5β1, α4β1, αvβ3 and a9b1 are not targets for the disintegrin alternagin-C from *Bothrops alternatus* venom (Souza *et al*., 2000). Cominetti *et al*. (2003) have recently shown that BaG, another metalloproteinase/disintegin from *Bothrops alternatus* venom, binds to α5β1 in K562 cells grown in suspension (as are HL60 cells). We found that the apoptotic index increased in the HL-60 cell line in a dose-dependent manner after treatmen twith whole venom and Balt-LAAO-II (Figure 7). Most of the biological effects of L-amino acid oxidases are believed to be due the secondary effect of H_2_O_2_ produced by enzymatic reactions (Massey and Curti, 1967; Tan and Ponnudurai, 1992). As H2O2 is permeable, it crosses the cellular plasmatic membrane, linking to DNA and causing cellular apoptosis. The authors suggests that this mechanism could be related to significant inhibition of in vitro cell growth HL-60 by action of L-amino acid oxidases Bal-LAAO-II. Furthermore, L-amino acid nutrients that are substrates of L-amino acid oxidases reactions are consumed in a way that can avoid proliferation of tumour cells (Kanzawa N et al., 2004) because they need the correct environment in which to develop and proliferate.

These results demonstrate that *Bothrops alternatus* venom contains an important substance the future anticancer drug development from snake venom and its mechanism of action. Lamino acid oxidases could effectively trigger cell apoptosis in vitro. Further studies will be necessary to be carried out to discover the structure and the sites of activity of Balt-LAAO-II.

## Acknowledgments

The authors thank José Ilton dos Santos and Gilberto Franchi Jr. for technical assistance.

## Captions figures

**Table 1.**
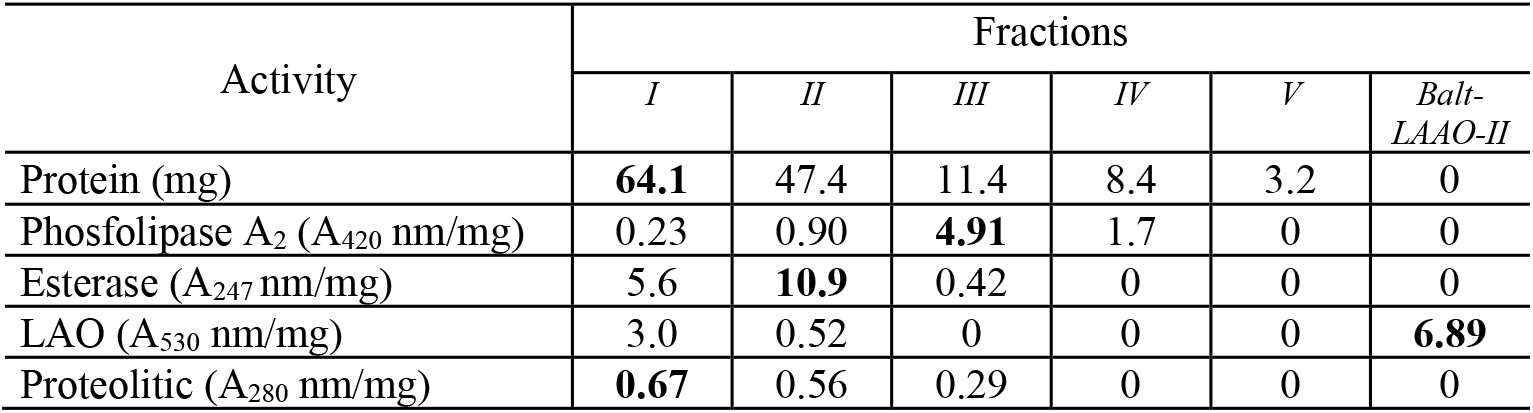
Protein content and enzymatic activities of fractions obtained by molecular exclusion of *Bothrops alternatus* snake venom and Balt-LAAO-II. Bold values indicates highest activities fractions.

| Activity | Fractions |  |  |  |  |  |
| --- | --- | --- | --- | --- | --- | --- |
|  | *I* | *II* | *III* | *IV* | *V* | *Balt-LAAO-II* |
| Protein (mg) | **64.1** | 47.4 | 11.4 | 8.4 | 3.2 | 0 |
| Phosfolipase A<sub>2</sub> (A<sub>420</sub> nm/mg) | 0.23 | 0.90 | **4.91** | 1.7 | 0 | 0 |
| Esterase (A<sub>247</sub> nm/mg) | 5.6 | **10.9** | 0.42 | 0 | 0 | 0 |
| LAO (A<sub>530</sub> nm/mg) | 3.0 | 0.52 | 0 | 0 | 0 | **6.89** |
| Proteolitic (A<sub>280</sub> nm/mg) | **0.67** | 0.56 | 0.29 | 0 | 0 | 0 |

